# Cryopreservation of Human Cortical Organoids Using Vitrification

**DOI:** 10.1101/2025.04.08.647634

**Authors:** Sandra Mojica-Perez, Kyle Stokes, Samantha Jacobs, Joy Huang, Shivanshi Vaid, Yukun Yuan, Caroline A. Pearson, Daniel Montes, Andrew Tidball, Debora VanHeyningen, Jonathan Gaillard, Louis T. Dang, M. Elizabeth Ross, Lori L. Isom, Michael Uhler, Wei Niu, Jack M. Parent

## Abstract

Cryopreservation at ultra-low temperatures is a valuable tool for preserving cells and tissues used in research. However, few protocols exist for the preservation of brain organoid models. Current methods for preserving human cortical organoids (hCOs) rely on conventional slow cooling approaches with organoids suspended in a medium containing a cocktail of cryoprotectants. In contrast, we have optimized a vitrification technique previously used to cryopreserve human embryos and oocytes for application to hCOs. We have successfully cryopreserved hCOs that were generated by two different protocols. The vitrified organoids demonstrate a growth rate, cytoarchitecture, cell type composition and electrical activity comparable to non-vitrified controls. Our hCO cryopreservation method provides a useful alternative approach for bio-banking and cross-institutional collaboration using cortical organoids as their model system.

**Highlights:**

- Human pluripotent stem cell (hPSC) derived hCOs can be cryopreserved using vitrification.
- Vitrified hCOs develop similarly to non-vitrified controls.
- Vitrified hCOs can be stored long-term and shipped to other institutions.

**Motivation:** Many methods of cryopreservation have been developed for the maintenance of cell lines, organoid models and tissues. Most organoid models have been successfully preserved using conventional slow-cooling methods; however, human cortical organoids have been difficult to preserve. Our findings demonstrate that the vitrification of cortical organoids preserves their structure, cell type diversity and function upon rewarming and continued culture.

## Introduction

Human brain organoid models are self-organizing three-dimensional (3-D) culture systems that recapitulate many aspects of brain development and function. They have become an invaluable tool for understanding human biology. These culture systems have a higher cellular diversity than two-dimensional (2-D) cultures and allow for more complex cell-cell interactions^1,2^.

Organoid systems have been widely used to study organ/tissue development, disease modeling, and to evaluate therapeutic efficacy. However, some obstacles with organoid culture methods make it challenging to apply these models for research. First, media and growth factors used during early differentiation steps are expensive. Second, brain organoid cultures may take several months or longer to reach the desired cellular diversity and neuronal maturity required for many applications. Third, unexpected termination of organoid experiments due to contamination or equipment malfunction can force investigators to lose valuable time and resources starting over. Fourth, intermural collaborations are complicated by large organoid batch variation between institutions. The ability to cryopreserve and then reconstitute organoids would enable investigators to preserve or share organoids from the same batch to mitigate these barriers.

Conventional cryopreservation methods utilizes a freezing medium containing 5-10% Dimethyl-Sulfoxide (DMSO) slow cooling (1°C/min) in a -80°C freezer overnight^3^. Cryogenic vials are then transferred to liquid nitrogen (LN_2_) at -195°C for long-term storage. This slow-cooling cryopreservation approach works well for several organoid systems, including gastrointestinal, lung, liver, and retina^4–7^. However, this method is unsuitable for cortical organoids and human brain tissue. Recently, a new cryopreservation medium with improved cryoprotective ability was developed, which composes of methylcellulose, ethylene glycol, DMSO and the Rho Kinase inhibitor Y-27632 (MEDY) which allows for the preservation of cortical organoids and brain tissue^8^.

Cryoprotectants (CPAs) such as the ones listed above are used to limit ice crystal formation. Due to the high water content in the cryopreservation medium, ice crystal formation is inevitable using the slow freezing approach^9^. Both the formation of ice crystals during freezing and ice melting during thawing disrupt intra- and extra-cellular structures, resulting in cell damage or death^10–12^. Adjusting concentrations of CPAs together with rapid warming is sufficient to prevent significant cell death in some organoid models^4–7,13^. To avoid the risks associated with the slow freezing method, we opted to utilize a vitrification strategy which has been shown to work well on very sensitive cell types like oocytes as well as whole embryos^14,15^.

The vitrification (turn into glass) approach to cryopreservation is the process of turning a liquid phase material into a solid phase through a sudden increase of viscosity which prevents ice crystallization^16^. The vitrification method of cryopreservation is most frequently used to cryopreserve oocytes and embryos for *in vitro* fertilization (IVF)^14,15^. Vitrification entails exposing cells or tissue to high concentrations of CPAs followed by a rapid cooling rate (∼2 – 5×10^6^°C/min), compared to the conventional method that cools at approximately 1-2°C/min^17^. Fast cooling causes the liquid in cells or tissues to solidify without crystallization. Warming after vitrification is also rapid (∼3×10^3^°C/min) to prevent ice formation followed by several washing steps to avoid CPA toxicity and osmolarity shock. To date, vitrification has been used on five 3D cell culture systems: intestinal^18^, lung cancer^13^, testicular^19^, mesenchymal stem cell spheroids^20^, and neuroectoderm spheroids^21^ but has not yet been shown to work on brain organoids. Vitrification and rewarming applied to whole rat kidneys and liver has been shown to preserve organ function after transplantation^22,23^. Taken together, the above research indicates that vitrification is an effective method for preserving tissues with complex structures.

In this study, we describe a vitrification approach that allows for the cryopreservation and long-term storage of human cortical organoids (hCOs). We find that hCOs preserved using this approach develop similarly to non-cryopreserved organoids. We also show that this method is suitable for storing hCOs long-term and shipping of samples between institutions.

## Results

### Vitrification is capable of cryogenically preserving human cortical organoids

To test whether vitrification would successfully preserve brain organoids, we chose to vitrify hCOs at day 20 post-differentiation (p.d.) to allow the hCOs to fully self-organize while still being small enough to facilitate easy penetration of the CPAs. We patterned human iPSC lines into “dorsalized” hCOs using a self-organizing single-rosette cortical organoid (SOSR-CO) protocol^24^. On day 20, the organoids (diameter of ∼0.5 mm) were separated into two groups: one underwent cryopreservation using our vitrification protocol (referred to as Vitri), which entails treating the organoids with increasing concentrations of the cryo-preservatives sucrose, DMSO, and ethylene glycol followed by immediate exposure to -195°C (Fig. 1; 2A, detailed protocol 1-11). The other group remained in culture (referred to as Control). One day post-vitrification (p.v.), hCOs were warmed through immediate exposure to 37°C rewarming solution (RS) followed by washes in decreasing levels of CPAs (Fig. 1; 2A; Detailed Protocol steps 12-21) and placed back into their usual medium alongside age-matched controls. SOSR-COs were analyzed for growth and cell composition at two timepoints (Fig. 2A).

**Figure 1.**
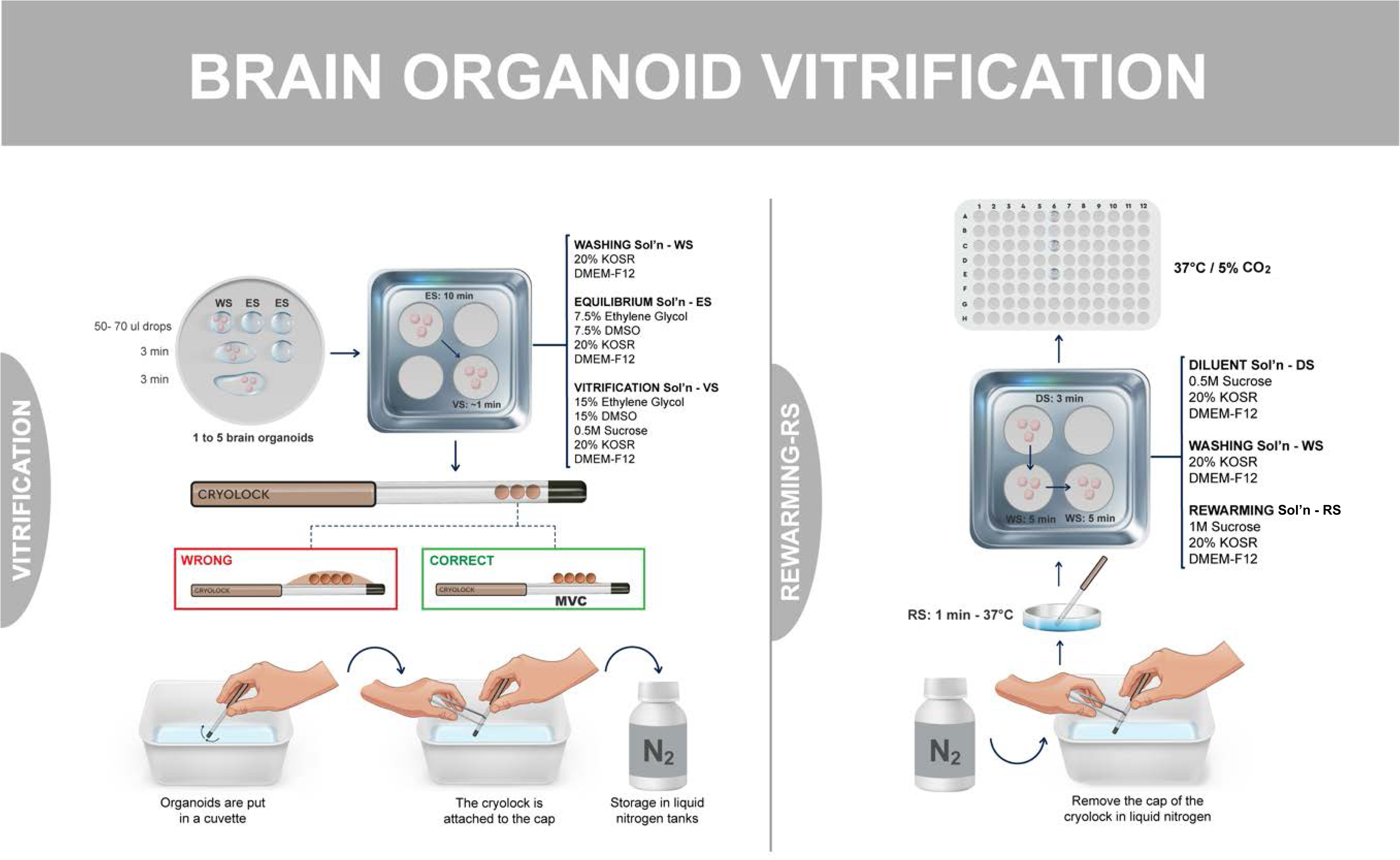
Overview of the vitrification protocol. Left: hCOs are vitrified by sequential placement in a series of solutions containing increasing levels of CPAs. hCOs are then placed in a Cryolock^®^ and all the CPA is removed followed by immediate immersion in LN_2_ for minimum volume cooling. These vitrified hCOs are then transferred to a cryo-storage tank for long-term storage. Right: Rewarming of hCOs requires immediate immersion of hCOs into a 37°C rewarming solution followed by sequential removal of CPAs.

**Figure 2.**
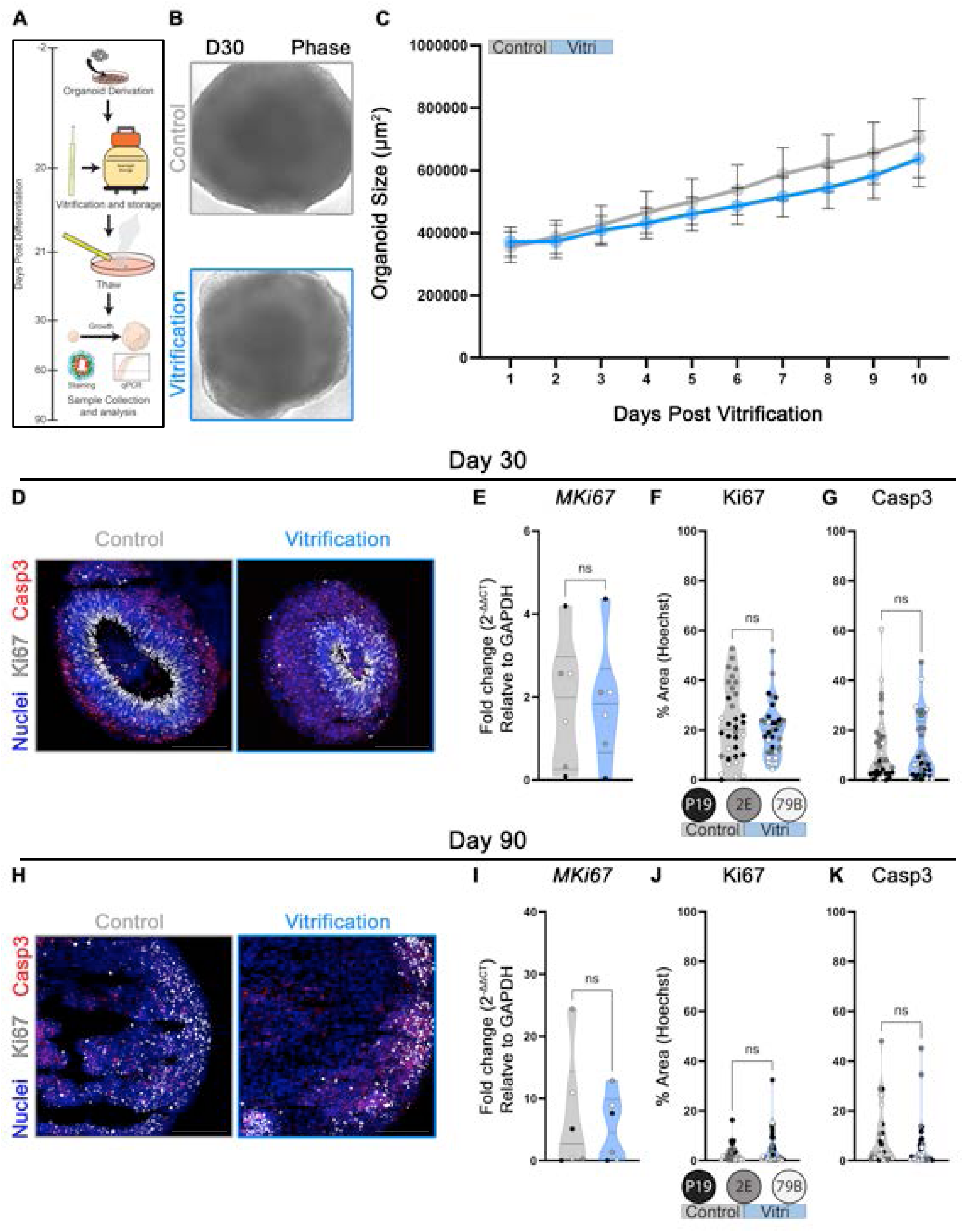
Vitrification does not impact organoid growth, cell proliferation or cell death. **A.** Schematic depicting workflow where SOSR-COs derived from 3 different human pluripotent stem cell (hPSC) lines (2E, gray; 79B, white; P19, black – see panel C) were grown to day 20. Half were vitrified and rewarmed the next day, and half remained in culture. Control and vitrified (Vitri) SOSR-COs were then collected and analyzed at Day 30 and 90 p.d. **B.** Representative phase images of control and Vitri SOSR-COs at 10 days post vitrification (day 30 p.d.). **C.** Control (gray) and Vitri (blue) SOSR-CO growth (µm^2^) was monitored for 10 days, beginning on the day of rewarming. No significant difference was seen in SOSR-CO growth between groups (p = 0.67, F_(1,10)_ = 0.19). Error Bars Represent SEM. **D.** Representative images of Day 30 p.d. control and Vitri SOSR-COs immunolabeled for MKI67 (Gray), Cas3 (Red), and Hoechst (Blue). **E-G.** No differences were observed between groups for the cell proliferation marker MKI67 quantified using RT-qPCR (**E**; p = 0.99) and IF (**F**; p = 0.49), or for Casp3 IF (p = 0.79). SOSR-COs from three separate hPSC lines (P19, black; 2E, gray; and 79B white) were measured. **H-K.** Proliferation (MKI67) and cell death (Casp3) were evaluated in Control (gray) and Vitri (blue) SOSR-COs at 90 days p.d. **H.** Representative confocal images of MKI67 (gray) and Casp3 (red) immunolabeled hCOs. MKI67 was not significantly different between groups using RT-qRT-PCR (**I**; p = 0.48) and IF (**J**; p = 0.29), and Casp3 IF was also unchanged (**K**; p = 0.19). Statistical tests: mixed effect analysis (**C**); Unpaired *t*-Test (**E-G, I-K**). Scale bars: 200µm.

For the first 10 days post vitrification (p.v.) (day 21-30 p.d.), the size of the SOSR-COs was monitored. We found that rewarming hCOs after vitrification did not cause any growth deficit (Fig. 2B-C). To assess differences in proliferation and apoptosis, we used immunofluorescence (IF) quantifications of MKI67 and cleaved caspase 3 (Casp3), respectively, with Fiji based on a previously described protocol^25^. We also used qRT-PCR to measure MKI67 transcript levels. At day 30 p.d. (day 10 p.v.) we found no significant differences in cell proliferation (MKI67^+^) or apoptosis (Casp3^+^) between vitrified and control SOSR-COs (Fig. 2D-G). At day 90 p.d. (70-days p.v.), Vitri SOSR-COs maintained the same levels of cell proliferation (MKI67^+^) and apoptosis (Casp3^+^) as controls (Fig. 2H-K).

We also assessed whether vitrification impacted the cellular composition of vitrified SOSR-COs. At day 30 we did not observe any significant difference in the number of neural stem/progenitor cells (SOX2^+^) or deep layer (CTIP2/BCL11B ^+^) neurons between vitrified and control SOSR-COs (Fig. 3A-E). We also found an equivalent number of intermediate progenitor cells (IPCs) marked by TBR2/EOMES^+^ and HOPX^+^ outer radial glia (oRG; Fig. 3F-J). Neurons (TUBB3/TuJ1^+^) were also found to be equivalent between vitrified and control SOSR-COs (Fig. 3K-M). These data indicate that vitrification of SOSR-COs at day 20 p.d. does not impact progenitor or neuronal cell numbers in day 30 p.d. hCOs.

**Figure 3.**
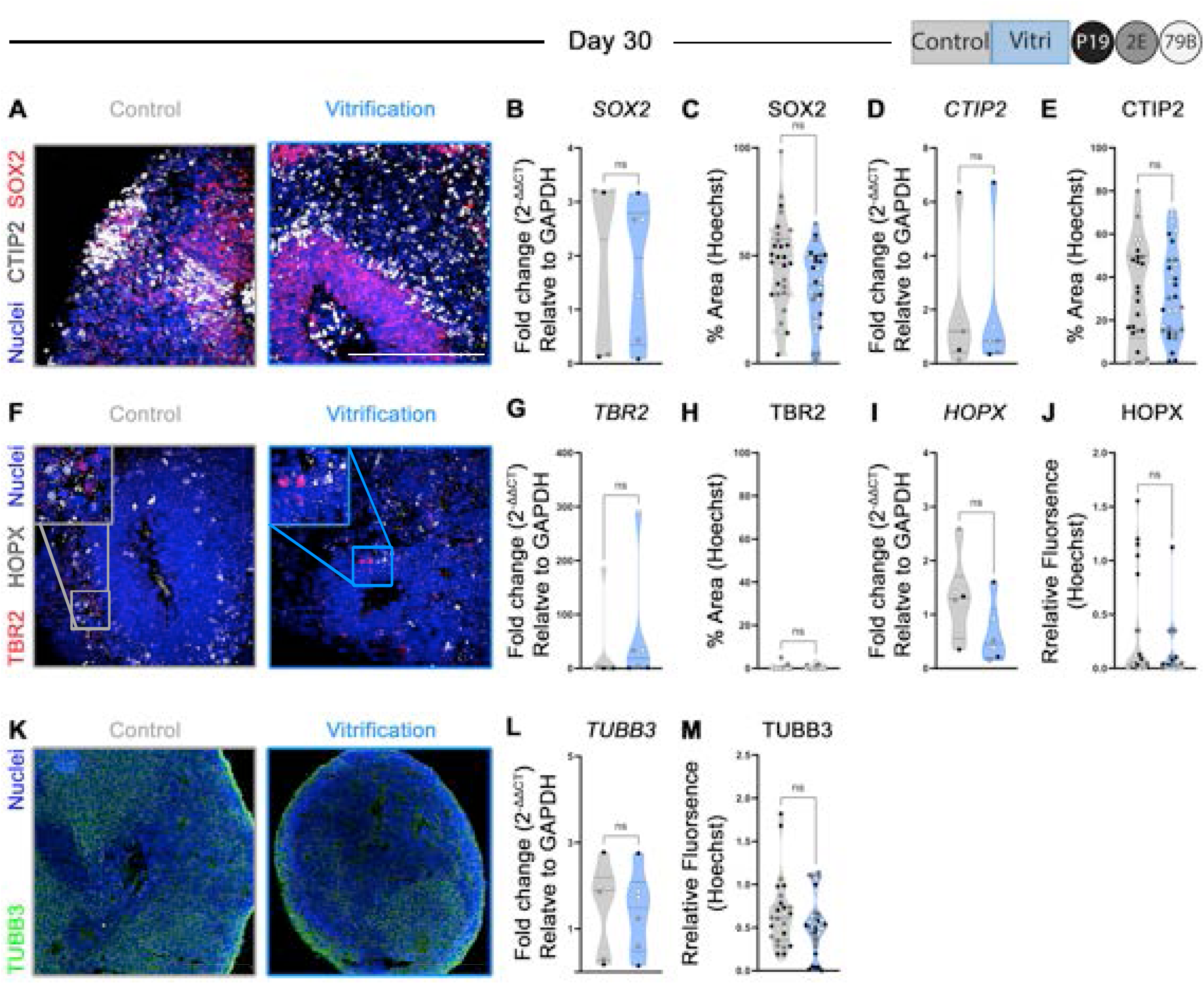
Day 30 SOSR-COs at 10 days post-vitrification have similar cellular composition to unvitrified organoids. **A.** Representative confocal images of control (left) and Vitri (right) organoids immunolabeled for the neural stem/progenitor cell marker SOX2 (red) and deep layer neuron marker CTIP2 (gray). **B-C.** SOX2 expression measured by RT-qPCR (**B**; p = 0.84) or IF (**C**; p = 0.07) did not differ between Vitri (blue) and control (gray) groups. **D-E.** Similarly, no changes were found in CTIP2 expression (**D**; p = 0.61) or IF (**E**; p = 0.92). **F.** Representative confocal images of control and Vitri SOSR-COs immunolabeled for IPC and oRG cell markers TBR2 (red) and HOPX (gray), respectively. **G-J.** Both TBR2 and HOPX expression are equivalent between groups by RT-qPCR (TBR2 in **G**; p = 0.61; HOPX in **I**; p = 0.14) and IF (TBR2 in **H**, p = 0.79; HOPX in **J**, p = 0.19). **K.** Representative confocal images of hCOs immunolabeled for the neuronal marker TUBB3 (green). **L**-**M.** Beta-III-tubulin expression (TUBB3) by RT-qPCR (**L**; p = 0.97) and IF (**M**; p = 0.2042) are not significantly different between controls (gray) and Vitri (blue) groups at day 30. Hoechst staining of nuclei in blue for panels A, F, K. The 3 different human pluripotent stem cell (hPSC) lines are shown in the plots as different colored dots (2E, gray; 79B, white; P19, black). Statistics: unpaired *t-*Test. Scale bars: 200µm.

To evaluate the potential effect of vitrification on maturity at a later stage of hCO development, we examined day 90 p.d. SOSR-COs for cell type diversity in a similar manner. Day 90 p.d. vitrified SOSR-COs showed no difference in transcript levels and a modest but statistically significant increase in immunolabeling for the neural stem/progenitor cell marker SOX2 (Fig. 4A-C). However, the levels of CTIP2^+^, SATB2^+^, and TUBB3^+^ neuronal populations (Fig. 4A, D-G, M-O), as well as the number of TBR2^+^ and HOPX^+^ neural progenitors (Fig. 4H-L), remained equivalent. The number and expression of GFAP^+^ labeled astrocytes also was not different between groups (Fig. 4P-R). Despite most cell types being at equivalent numbers, the increase in SOX2^+^ cells may indicate a slight delay in maturation in the vitrified organoids compared to age matched controls, but this remains to be definitively determined.

**Figure 4:**
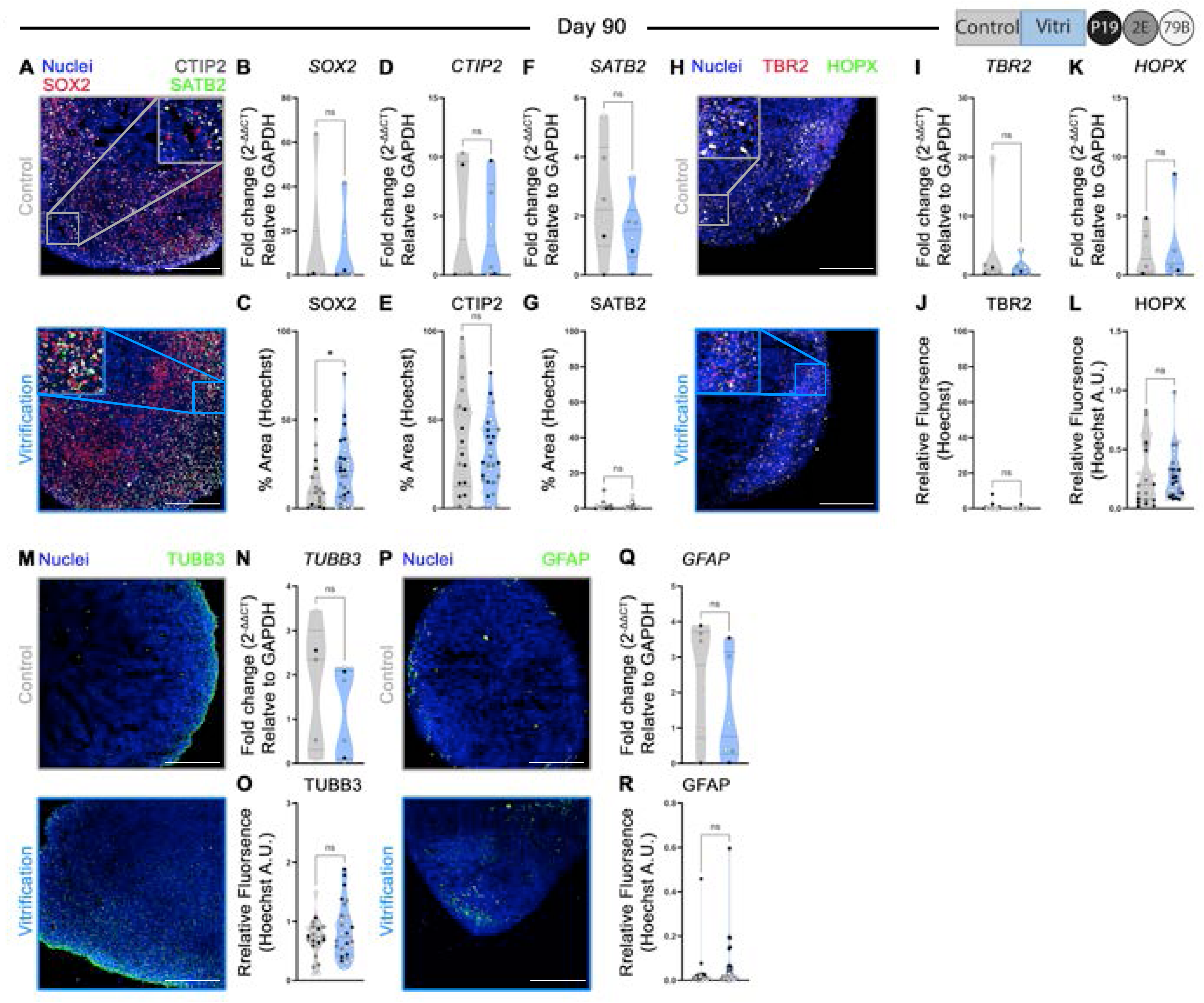
Vitrified SOSR-COs develop similarly to unvitrified controls. Day 90 p.d. SOSR-COs were assessed for cell type diversity. **A.** Representative confocal images of control and Vitri (bottom) organoids immunolabeled for the neural stem/progenitor cell marker SOX2 (red), deep layer neuron marker CTIP2 (gray), and superficial layer neuron marker SATB2 (green). Hoechst nuclear stain is in blue. **B-C.** SOX2 shows no difference in expression between control (gray) and Vitri (blue) groups by RT-qPCR (**B**; p = 0.13), but Vitri SOSR-COs have significantly more SOX2^+^ cells by IF (**C**; p = 0.03). **D-E.** CTIP2 expression by RT-qPCR (**D**; p = 0.47) and IF (**E**; p = 0.23) are unaffected by vitrification. **F-G.** Levels of SATB2 by RT-qPCR (**F**; p = 0.06) and IF (**G**; p = 0.89) do not differ between groups. **H.** Representative confocal images of control and Vitri SOSR-COs immunolabeled for TBR2 (red) and HOPX (green), with Hoechst nuclear stain (blue). **I-L.** The expression of both TBR2 (**I**, RT-qPCR, p = 0.55; **J**, IF, p = 0.19) and HOPX (**K**, RT-qPCR, p = 0.20; **L**, IF, p = 0.72) does not differ significantly between groups. **M.** Representative confocal images of the neuronal marker TUBB3. **N.** TUBB3 expression by qRT-PCR is unchanged (p = 0.26). **O.** TUBB3 IF intensity (p = 0.29) does not differ between controls (gray) and Vitri (blue) SOSR-COs at day 90. **P.** Representative confocal images of the astrocyte marker GFAP. **Q-R.** No significant differences are found between the mRNA expression or IF intensity of GFAP^+^ cells between vitrified and control samples (**Q**, RT-qPCR: p = 0.23; **R**, IF: p = 0.09). Statistics: unpaired *t-*Test (**B**-**G**, **I**-**L, N, O**, **Q**, **R**). * = p < 0.05. Scale bar: 200µm.

In addition to developing a structural and cellular composition that largely recapitulates the developing human cortex, hCOs are also capable of exhibiting neuronal and network activity^26–30^. We assessed activity by plating day 90 p.d. SOSR-COs onto MEAs, slowly transitioning them to BrainPhys medium and then recording spontaneous field potentials (Fig. 5A). Between weeks 18-21 we observed an equivalent number of active electrodes and similar levels of firing over a 5-minute recording between Vitri and control SOSR-COs (Fig. 5B, C). We continued to maintain SOSR-COs in BrainPhys medium until day 200 p.d. and performed whole-cell patch clamp recordings from organoid slices. We found no substantial differences in evoked action potentials (AP) from neurons in control and Vitri SOSR-COs (Fig. 5D). Input-output curves of evoked APs showed no significant difference between groups (Fig. 5E). Spontaneous excitatory postsynaptic currents (sEPSC) recordings also show comparable amplitude and frequency between control and Vitri neurons (Fig. 5F-J), indicating that vitrification does not prevent the formation of synapses in hCOs.

**Figure 5:**
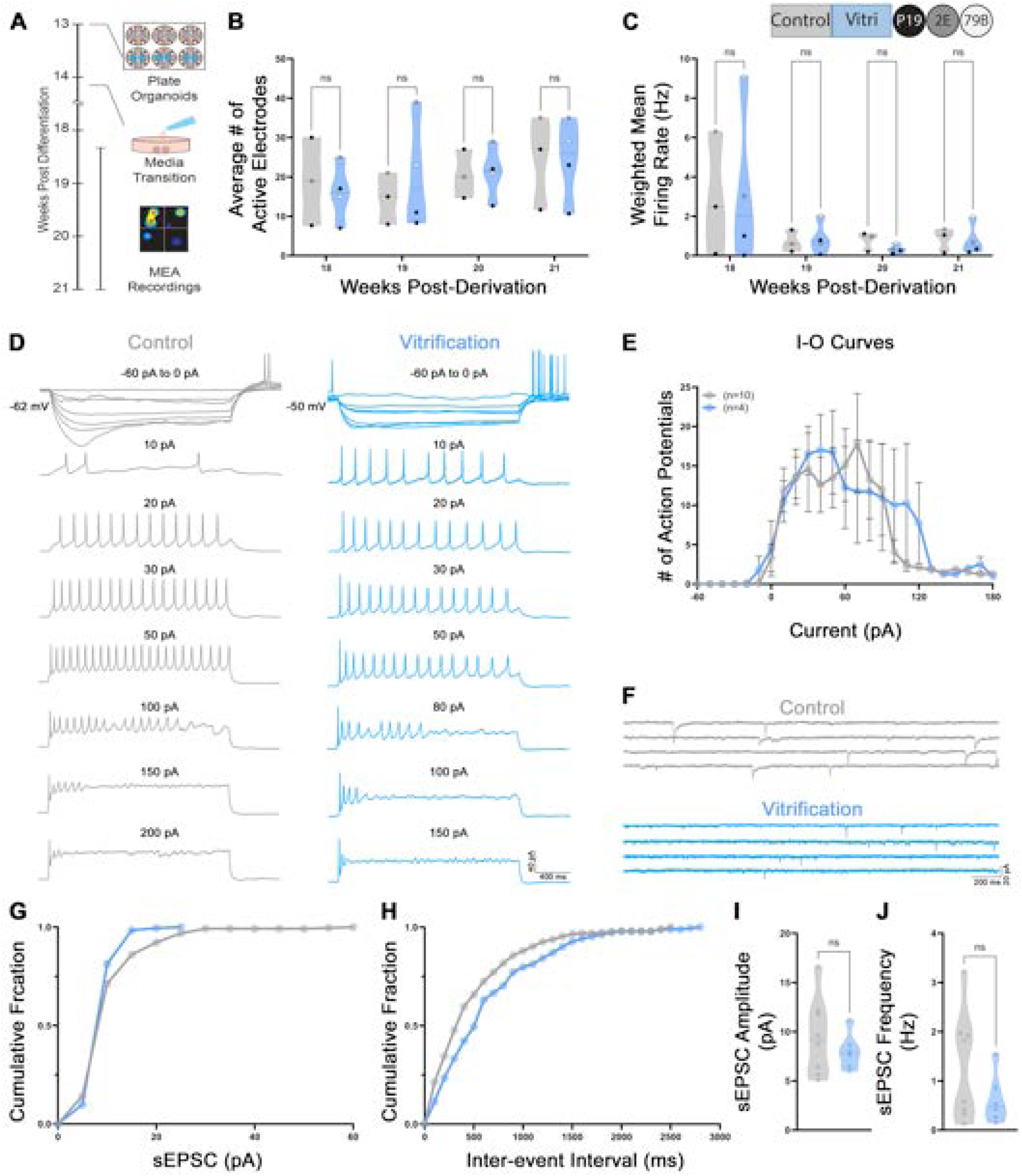
**A.** Schematic showing the protocol for hCO recordings on 6-well MEA plates. **B, C.** Weeks 18-21 SOSR-COs show equivalent numbers of active electrodes (**B**, F_(3, 20)_ = 0.23, p = 0.87) and spontaneous field potentials displayed as weighted mean firing rate in Hz (**C**, (F_3, 20_) = 0.47, p = 0.99) between control (gray) and Vitri (blue) organoids. **D.** Representative traces of evoked APs recorded by whole-cell patch clamp. **E.** Input-output (I-O) curves show similar levels of activity in neurons from control (gray, n = 10) and Vitri groups (blue, n = 4; F_(31, 379)_, p = 0.99). **F.** Representative traces depicting similar levels of sEPSCs in voltage-clamp recordings of hCO slices made from the two groups **G, H.** Graphs depict the cumulative fraction of spontaneous excitatory postsynaptic current (sEPSC) amplitude (pA) and inter-event interval (ms). **I, J.** Both sEPSC amplitude (pA; p = 0.4) and frequency (Hz; p = 0.19) are equivalent between control (gray, n = 8) and vitrified (blue; n = 6) neurons within day 200 SOSR-COs. Statistics: 2-way ANOVA with Šídák post-test (**B**, **C, E**); *t*-test (**I**, **J**).

### Vitrification of hCOs derived using a multi-rosette hCO protocol

Our SOSR-CO hCO protocol tends to make smaller cortical organoids due to starting with a single rosette. To test whether our vitrification protocol works on a commonly used multi-rosette hCO (MR-hCOs) derivation method that generates larger organoids, we applied vitrification to an hCO protocol developed by Sergiu Paşca’s group^31^ that is widely used by investigators. MR-hCOs were vitrified on day 20 as per our previous experiment (Figs. 1, 2A). As expected, the MR-hCOs were significantly larger than SOSR-COs at the same timepoint and thus the CPA cocktail had a larger volume to penetrate. MR-hCOs were rewarmed the day after vitrification and maintained in culture to day 90 p.d. (day 70 p.v.). Similar to our previous results, the MR-hCOs at day 90 p.d. did not show any differences in size (Fig. 6A, B), MKI67^+^ proliferating cells, or Casp3^+^ apoptotic cells (Fig. 6C, H, I), nor did we observe any difference in the abundance of SOX2^+^ neural stem/progenitor cells, TBR2^+^ IPCs, HOPX^+^ oRG or TUBB3^+^ neurons (Fig. 6D-G, J-O). However, we did find a significant decrease in the number of astrocytes labeled by IF for GFAP in Vitri organoids (Fig. 6G, P). These data demonstrate that our vitrification protocol works with both single- and multi-rosette hCO protocols with minimal impact on the development of the hCOs.

**Figure 6.**
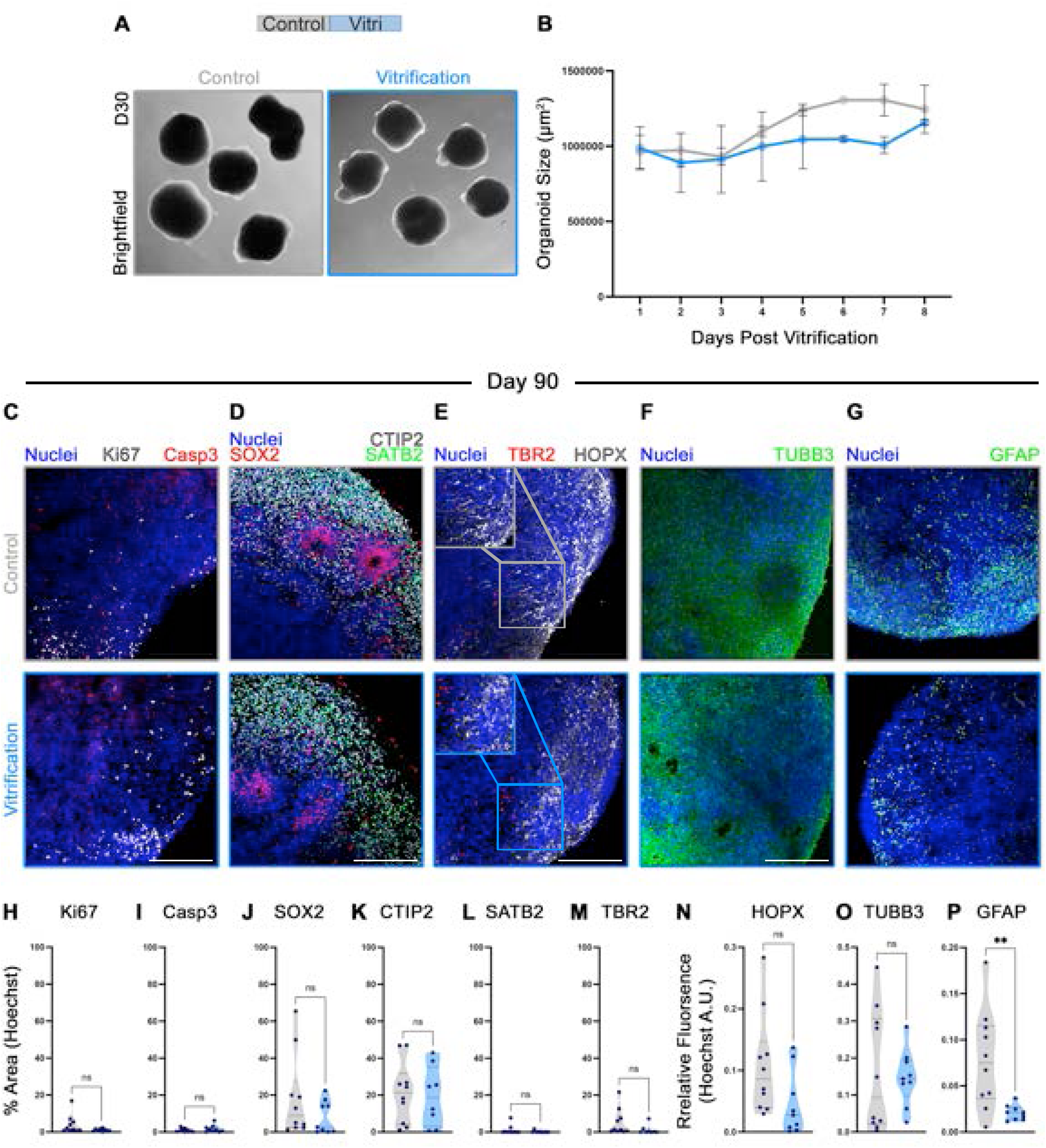
Vitrification can be applied to multi-rosette cortical organoid protocols. **A.** Representative brightfield images of control (gray) and Vitri (blue) hCOs generated using the multi-rosette protocol (MR-hCOs) at day 30 p.d. (10 days p.v.). **B.** Vitri (blue) MR-hCOs maintained a similar growth rate as controls (gray; p = 0.07, F_(1,16)_ = 3.84). Error bars represent SEM. **C-G.** 90-day p.d. (day 70 p.v.) P-hCOs were stained for selected cell type markers. Representative confocal image of control and Vitri MR-hCOs that were immunolabeled with antibodies to **C**: KI67 (gray) and Casp3 (red); **D**: SOX2 (red), CTIP2 (gray) and SATB2 (green); **E**: TBR2 (red) and HOPX (gray); **F**: TUBB3 (green); and **G**: GFAP (green). Hoechst nuclear counterstains are in blue. **H-P.** Vitri (blue) P-hCOs have the same number of MKI67^+^ (**H**; p = 0.11;), Casp3^+^ (**I**; p = 0.33), SOX2^+^ (**J**; p = 0.22), CTIP2^+^ (**K**; p = 0.80), SATB2^+^ (**L**; p = 0.32), TBR2^+^ (**M**; p = 0.15), HOPX^+^ (**N**; p = 0.06) and TUBB3^+^ (**O**; p = 0.81) cells, but show a reduced number of GFAP^+^ cells (**P**; p = 0.006) compared to unvitrified controls (gray). Statistics: two-way ANOVA (**B**), unpaired *t-*Test (**H-P**). Scale bars: 200µm.

### Vitrification of hCOs allows for long-term storage and shipment to other institutions

To determine whether vitrified hCOs can be stored over a longer period, we rewarmed SOSR-COs that were vitrified on day 35 p.d. and stored in LN_2_ for over 1 year. After rewarming, SOSR-COs were collected at day 60 p.d. and compared to non-vitrified control hCOs derived from the same experiment. The vitrified SOSR-COs demonstrated a significant increase in MKI67^+^ and Casp3^+^ cells, indicating a possible regenerative response (Fig. 7A-C). We also found an increased number of CTIP2^+^ deep layer neurons without changes in the relatively low number of SATB2^+^ superficial layer neurons expected at this timepoint (Fig. 7D-F). No significant differences in SOX2^+^ neural stem/progenitor cells or TUBB3^+^ neurons were observed (Fig. 7G-I). Thus, SOSR-COs stored for at least 1 year survived but displayed some cell type-specific differences at day 60 p.d. These differences may reflect the effects of longer-term storage that would need to be addressed, or they instead could result from performing the vitrification at a slightly later developmental timepoint (day 35 instead of day 20).

**Figure 7:**
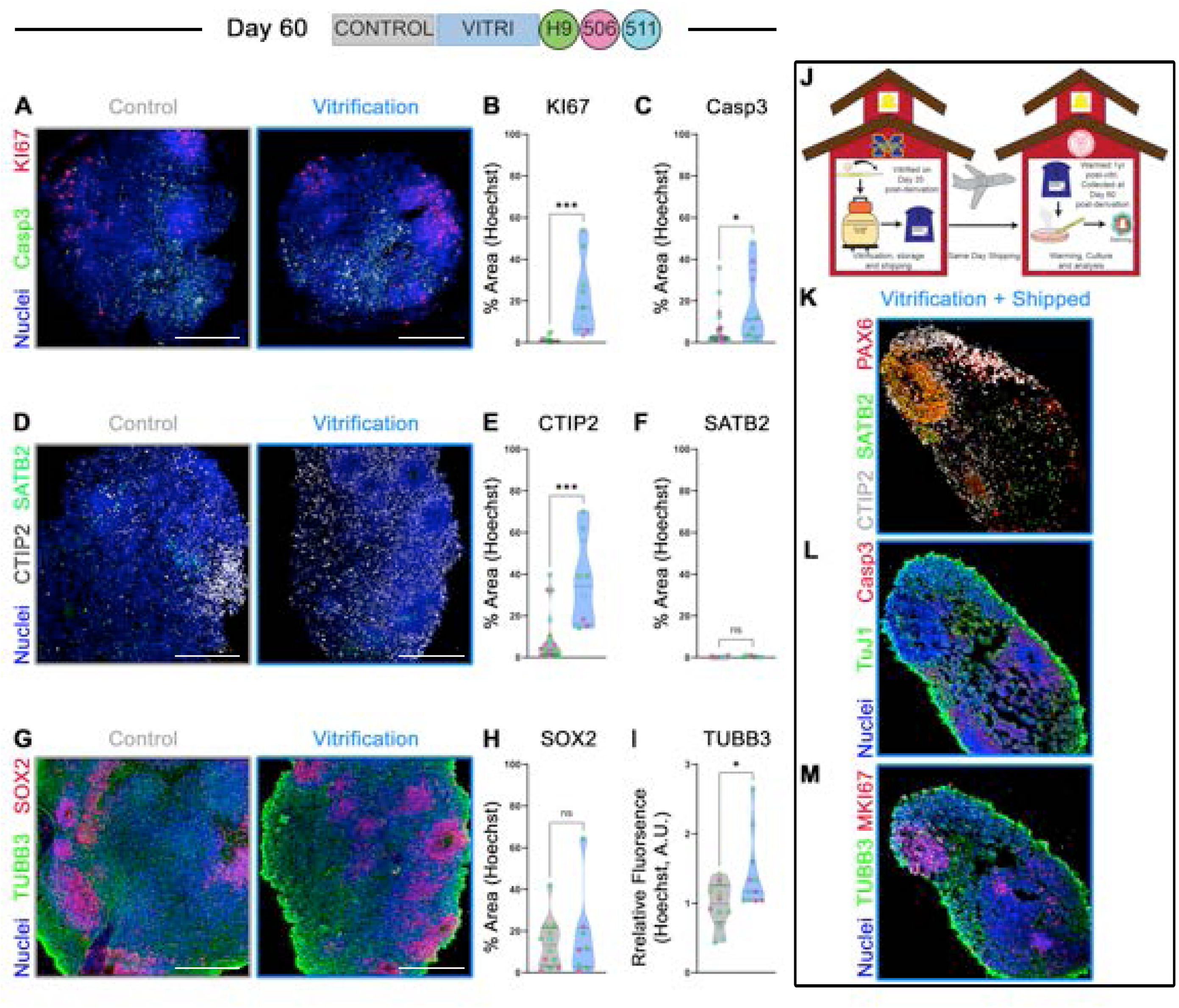
Vitrified SOSR-COs can be stored long-term and shipped to other labs for reconstitution. **A–H.** Representative confocal images of control and Vitri SOSR-COs. The Vitro group were vitrified on day 35 p.d., stored for over 1 year in LN_2_, rewarmed and cultured until day 60 p.d. Significant increases were found in KI67^+^ (**A**, **B**; red in **A**; *t*-Test, p = 0.0005), Casp3^+^ (**A**, **C**; Green in **A**; *t*-Test, p = 0.04) and CTIP2^+^ (**D**, **E**; gray in **D**; *t*-Test, p = 0.0002) cells in the Vitri (blue) group compared to controls (gray). No significant differences were observed in SATB2^+^ (**D**, **F**; green in **D**; *t*-Test, p = 0.06), SOX2^+^ (**G**, **H**; red in **G**; *t*-Test, p = 0.81) or TUBB3^+^ (**G**, **I**; green in **G**; *t*-Test, p = 0.11) cells between Vitri (blue) and control (gray) SOSR-COs. **J.** Graphical representation of protocol used to ship vitrified SOSR-COs from the University of Michigan to Weill-Cornell Medical Center. **K-M.** Representative images of SOSR-COs shipped to Cornell after freezing at day 35 p.d. and storage for > 1 year, then re-warmed and cultured to day 60 p.d. (day 25 p.v.). The SOSR-COs showed immunoreactivity for PAX6 (red in **K**), CTIP2 (gray in **K**), SATB2 (green in **K**) and TUBB3/TuJ1 (green in **L**, **M**). They also showed expected amounts of Casp3 (red in **L**), and KI67 (red in **M**). Hoechst nuclear counterstains are in blue. Unpaired t-Test (**B, C, E, F, H, I**). * = p ≤ 0.05; *** = p ≤ 0.0005. Scale bar: 200µm.

We next tested whether our cryopreserved SOSR-COs could maintain their viability and structure after long-distance shipping. We vitrified SOSR-COs on day 35, stored them for over a year, and then shipped them from the University of Michigan to our collaborators at Weill Cornell Medical School in New York City using overnight delivery (Fig. 7J). Our collaborators, new to the vitrification protocol, were then instructed on how to rewarm the organoids using our method. They then cultured these SOSR-COs to day 60 p.d. (25 days p.v.). Although there were too few SOSR-COs to quantify cell type diversity, we observed that the SOSR-COs survived the long-term storage and long-distance shipping and expressed the expected makers found at the 60-day timepoint, including PAX6, CTIP2 and SATB2 (Fig. 7K). The SOSR-COs also displayed the expected levels of Casp3 and MKI67 expression and abundant TUBB3^+^ neurons (Fig. 7L, M). These data indicate that SOSR-COs survive long-term cryopreservation and long-distance shipping.

## Discussion

The use of vitrification for cryopreserving many different cells and tissues has increased in recent years^20^. Previous studies have shown that 2D and 3D cell culture models are amenable to vitrification protocols^4,13,19,20,22,32^. Even whole tissues with complex architecture including rat liver and kidney were vitrified and, upon rewarming, showed preserved organ function after transplantation into a host^22,23^. Although brain organoids have been studied for well over a decade, the ability to reliably preserve them has been challenging^21,26^. In this study we have demonstrated that hPSC-derived hCOs, generated using two separate protocols at 3 weeks p.d., can be reliably preserved in LN_2_ and rewarmed with limited impact on organoid development and function.

We find that hCOs undergoing vitrification survive the rewarming process and show equivalent levels of cell division, cell death and cell type-specific markers compared to unvitrified controls. Additionally, we show that neurons from hCOs that have been vitrified, rewarmed and cultured to relative maturity demonstrate equivalent levels of neuronal activity compared to non-vitrified controls, indicating that the functionality of vitrified hCOs is retained. Our findings also show that this protocol may be applied successfully to a commonly used multi-rosette hCO derivation protocol, suggesting that it will be useful for many neural based organoid systems. Finally, we show that our protocol can be used for the long-term storage of hCOs and that vitrified hCOs can be shipped between institutions with little detriment to the hCO, although we did identify some developmental changes induced by long-term storage.

Why long-term storage in LN_2_ altered the subsequent development of the hCOs is uncertain at present. This finding may reflect toxic or anti-oxidant effects of some CPAs including DMSO which was used in our protocol^33^ or may relate to the cryopreservation process in general, as a recently described hCO cryopreservation method also showed a significant increase in the progenitor cell zone with a slight but non-significant increase in SOX2^+^ cells.^8^ Alternatively, the changes after long-term storage in our studies may result from the timing of hCO vitrification as they were vitrified at day 35 p.d. instead of day 20 p.d. hCOs at the later time point contain more heterogenous and possibly more fragile cell populations.

Previous work has shown that different cell types have variable tolerances to CPA concentrations and osmotic pressure^34,35^. We do not believe this was related to time in storage or CPA penetration. Previous studies have observed that the length of time in cryopreservation had minimal effects on cells after rewarming^4,8,36–41^, and the day 20 MR-hCOs were much larger at the time of vitrification than day 35 SOSR-COs used in the long-term storage experiments, indicating that CPA penetration was not problematic.

A recent article published by Xue et al. describes a slow cooling method (MEDY) that was able to cryopreserve multiple neural based organoid models as well as human brain tissue.^8^ This work shows that neural tissues that have undergone a freeze/thaw can maintain their organization and function with minimal impact on cell type diversity and excitability. Our method, in comparison, has a few notable advantages. First, the amount of time spent in CPAs is reduced from hours to minutes and the time spent in CPAs did not need to be augmented based on hCO size. Our protocol also does not require any supplemental culturing following rewarming, allowing hCOs to be immediately placed back in their native culture conditions. Second, vitrification prevents ice formation, and the 1-min rewarming step limits the potential shearing damage that may occur due to ice shifting which frequently occurs with slow freezing protocols^9^. Finally, the size and shape of the Cryolock® as well as its ability to hold 3-6 organoids each, minimizes the space requirements for long-term storage, whereas the MEDY protocol uses one cryotube per organoid which could take up a considerable amount of space depending upon the experiment.

In summary, we demonstrate here that vitrification is a viable alternative strategy for the cryopreservation of hCOs using two organoid derivation protocols. We have also shown that it is possible to store vitrified hCOs for over a year and ship them long-distance with minimal alterations in organoid growth and cellular diversity. This approach will be useful for sharing hCOs with labs that would like to perform research on organoids but do not have the necessary expertise with hPSC culture methods or cannot afford the financial barriers of long-term culture. Additionally, this protocol may allow for the creation of brain organoid biobanks to avoid sample loss and to allow for large-scale sharing of patient tissue.

### Limitations of the Study

Vitrification of hCOs can maintain cell diversity and neural activity comparable to unvitrified controls. With our analyses, however, we cannot exclude differences in less common cell types that were not examined. The protocol could also be refined as we only tested one CPA cocktail formulation and only performed vitrification at two timepoints (Day 20 and 35 p.d.). Thus, the protocol may be further optimized by exploring other time points or CPA cocktails. Finally, our protocol is limited in scalability due to the size of the Cryolock^®^, although we are working with the manufacture to adapt the Cryolock^®^ technology to allow for more organoids to be stored in the same Cryolock^®^.

## Acknowledgments

We would like to thank the IVF Store for providing us with samples of the Cryolock^®^ to perform pilot experiments. We would also like to extend a special thank-you to Vanessa Guzmán and Andrea Merizalde for generating the graphics for Figure 1.

## Author Contributions

S.P.M.P. conceptualized the methodology and developed the protocol. Experiments were conducted and data was analyzed by S.P.M.P., K.S., S.J., S.V., C.A.P., D.M., J.H. and J.G. Patch clamp recordings were performed by Y.Y. Shipped SOSR-COs were cultured and imaged by C.A.P. MR-hCOs were provided by D.V. Original draft written by K.S., S.P.M.P. All authors reviewed and edited the manuscript. W.N., J.M.P., M.U., M.E.R., L.D, L.I. contributed to experimental design and supervised different aspects of the study.

## Declaration of Generative AI is Scientific Writing

The authors did not use generative AI in the preparation of this manuscript.

## STAR Methods

### hPSC culture

All hPSC experiments were conducted following prior approval from the University of Michigan Human Pluripotent Stem Cell Research Oversight (HPSCRO) Committee. Male A2 (Supplemental Figure 1) and CON2E^42^ lines were reprogrammed to induced pluripotent stem cells (iPSCs) from human foreskin fibroblasts using a previously described protocol^42^. The female 79B iPSCs were reprogrammed from the blood of a healthy female that we purchased from ZenBio (#SER-WB10ML-SDS; Supplemental Figure 1). The P19^24^, 506, and 5ll cell lines were derived from the female WA09 (H9) human embryonic stem cell (hESC) line (WiCell, # WB68074) and subjected to CRISPR gene editing. The 511 line is a CRISPR edited line with a heterozygous mutation in *STXPB1* generated by using the Precision gRNA synthesis kit (Thermo, #A29377) and transfected with TrueCut™ Cas9 protein v2 (Thermo, #A36496) using the Neon™ transfection electroporation system (Thermo, #NEON1S). The sgRNA sequence is CGTAGACAAACTCTGCCGAG targeting *STXBP1.* The 506 line is an isogenic control line in which *STXBP1* is not edited despite exposure to CRISPR reagents. Single nucleotide polymorphism DNA microarray (SNP-Chip) was performed to confirm genomic integrity.

hPSC lines were maintained in feeder-free conditions and cultured on Geltrex-coated (1:100 dilution; Thermo, #A1413302) 6-well plates (Thermo, #11330032) in either mTeSR1, mTeSR™ Plus (STEMCELL Tech, #85850; #100-0276,) or StemFlex™ (Gibco, #A3349401) stem cell maintenance medium at 37°C with controlled humidity and 5% CO_2_. Cells were routinely passaged at 80% confluency by washing with phosphate buffered saline without calcium and magnesium (PBS; Gibco, #14190144), then lifted off the plate by incubating in 0.8mM ethylenediaminetetraacetic (EDTA) for 5 min diluted from AccuGENE^®^ 0.5M EDTA Solution (Lonza, #51201)^43^. Mycoplasma testing was carried out periodically using PCR amplification of ribosomal DNA of mycoplasma with the primer set (5′-TGCACCATCTGTCATTCTGTT-3′ and 5′-GGGAGCAAACAGGATTAGATA-3′).

### Organoid differentiation protocols

#### SOSR-CO differentiation

The self-organizing single rosette organoid (SOSR-CO) protocol was modified slightly from our published method^24^. Briefly, hPSC lines between passage 8-20 were passaged using StemPro™ Accutase™ (Thermo, #A1110501) and replated onto Geltrex-coated (1:50 dilution in DMEM/F12), 12-well plates at 6×10^5^-2×10^6^ cells/well (cell line dependent) in either mTeSR™ Plus or StemFlex™ with 10µM rho-kinase inhibitor (Y-27632; Tocris, #1254). Rho-Kinase inhibitor was removed the next day, and 2 mL of stem cell medium was replaced daily until cells reached 80-100% confluency. A full media change was done with modified 3N culture medium^44^ without vitamin A, supplemented with 4 inhibitors to facilitate differentiation down a neuronal path: 2µM DMH1 (Tocris, #4126), 2µM XAV939 (Cayman Chemical, #13596), and 10µM SB-431542 (Cayman Chemical, #13031) (Day 0). Daily 75% media changes with the addition of 1µM cyclopamine (Cayman Chemical, #11321) were performed on days 1-3. On day 4, the monolayer was cut into squares using the StemPro EZ passage tool (Fisher, #23181010), liberated using a hypertonic sodium citrate solution^45^ and washed with preconditioned 3N medium. Liberated squares were diluted with an additional 2 mL of fresh 3N culture medium containing 4 inhibitors (DMH1, XAV939, SB-431542, Cyclopamine). 200µL of resuspended squares were then transferred into each well of a 96-well plate preincubated at 37°C for 30min with 35µL of 100% Geltrex/well. On day 6, a half medium change was performed with 3N media containing 3µM CHIR99021 (Cayman, #13122). On Day 10 the organoids were removed using the STRIPPER^®^ Micropipetter with a bent 275µM tip (Cooper Surgical, #MXL3-STR; #MXL3-275) and plated individually in a low adherence U-bottom 96-well plate (Corning, #7007) with 200µL of 3N medium without vitamin A and with 3µM CHIR99021, 20ng/mL Brain Derived Neurotrophic Factor (BDNF) and 20ng/mL Neurotrophic factor-3 (NT3) (Peprotech, #450-02; #450-02). On day 12, half-media changes every-other-day were initiated with 3N with vitamin A, 20ng/mL BDNF and 20ng/mL NT3. From day 30 onward SOSR-COs were transferred to low-adherence 24-well plates (Corning, #3475) in 400µL 3N media with vitamin A. Media was exchanged every other day after day 30.

#### MR-hCO differentiation

This protocol was modified slightly from the previously published version^31^. Briefly, hPSCs were dissociated to single cells using Accutase, and 3×10^6^ cells were added to a single well of an Aggrewell 800 plate (STEMCELL Tech, #34811) with 2 mL of mTesR1 medium (STEMCELL Tech, #85850) containing 50µM Rho-Kinase inhibitor (Day -1). A full media change was performed the following day (Day 0) by replacing mTesR1 with Neural Induction Media [NIM; Neurobasal-A (Thermo, #A2477501), 20% Knockout serum replacement (Thermo, 10828028), 0.5% GlutaMAX 100X (Thermo, #35050061), 1% MEM NEAA (Thermo, #11140035), 1% Pen/Strep (Thermo, 15140122), 10µM β-mercaptoethanol (Thermo, 21985023)] containing 5µM Dorsomorphin (Tocris, #3093) and 10µM SB-431542. On day 1, embryoid bodies (EBs) were formed and transferred to a 100mm dish (Corning, #430591) with 10 mL NIM medium containing Dorsomorphin and SB-431542 and placed on a rotating shaker at 37°C with 5% CO_2_. Daily media changes were done until day 5. From days 6-14, media was exchanged daily with Neural Media [NM; Neurobasal-A, 1% GlutaMAX, 1% Pen/Strep, 2% B-27 Supplement -A (Thermo, #12587010)] containing 20ng/mL Epidermal Growth Factor (EGF; Millipore, #01-102) and 20ng/mL Fibroblast Growth Factor-2 (FGF2; R&D Biosystems, #233-FB). On day 15, organoids were removed from the shaker and transferred into a 6-well plate (10/well) to avoid overcrowding, and the media was exchanged every other day. On day 25, the media was switched to NM containing 20ng/µL BDNF and 20ng/µL NT3 and changed every other day. From day 43 onward the media was exchanged every fourth day with NM only.

### Cryopreservation and Rewarming Protocol

#### Vitrification and Rewarming

In a HERA Guard clean bench (Thermo) using a Stripper^®^ CC micropipetter with 600µm tip (Cooper Surgical, #MXL3-STR-CC; #MXL3-600), 3-6 (size dependent) day 20 p.d. hCOs were placed into a 70µL droplet of washing solution (WS) which was then merged with a droplet of equilibration solution (ES) and incubated for 3min at room temperature (RT). A second bead of ES was merged with the first two and incubated for another 3min. hCOs were then transferred to 1 well of a 4-well plate (Fisher, #14444) containing ES and incubated for 10 min at RT. hCOs were again transferred to a well continuing vitrification solution (VS) for 1min then immediately transferred to a Cryolock® (IVFStore, #SCL-R-CT-B) where all remaining media was removed. The Cryolock® was then submerged into liquid nitrogen (LN_2_) and capped while still submerged. The capped Cryolock® containing the hCOs was then transferred to a 13mm Goblet affixed to a Cryo-Cane (IVF store, #13-100-CLR, Thermo, #5015-0002) and placed in LN_2_ tank for long-term storage. For rewarming, an uncapped Cryolock® was submerged into 37 rewarming solution (RS) in a 35mm petri dish (Greiner, #627161) for 1min. hCOs were transferred using a cut-off p200 pipette to a well containing diluent solution (DS) and incubated for 3min at RT. We then performed two 5-min incubations in WS at RT and transferred hCOs into native media for continued culture. See supplemental table 1 for solution compositions. Detailed protocol descriptions can be found in Figure 1, supplemental videos 1 and 2, as well as the supplemental experimental procedures.

### Shipping Vitrified Organoids

For information on how to prepare the vapor shipper dewar (IVF Store, #10817330) and the protective shipping container (IVF Store, #20750409) for organoid shipment, please refer to the manufacturer’s <u>operation manual</u> and add canes containing Cryolocks^®^ right before shipping. The dewar should maintain its temperature for 21 days after removing the LN_2_. Overnight shipping arrangements should be made accordingly.

### Immunocytochemistry

The hCOs were fixed at day 30 with 4% paraformaldehyde (PFA; Fisher, #AA433689M) in PBS for 30min at 4°C, while on day 90 hCOs were fixed for 1h in 4% PFA at 4°C. Organoids were then washed 3 times for 10min in 1×PBS and incubated overnight in 30% sucrose in 1×PBS at 4°C. Organoids were embedded in O.C.T. (Thermo, #23-730-571) in Tissue-Tek Cryomolds (Sakura, #32663) and frozen on dry ice before being stored at -80°C. The blocks were sectioned on a cryostat (Leica) at 20µm thickness. After 2 h of drying at RT, the sections were outlined using a hydrophobic A-PAP Pen (Ted Pella, #22309). Once the ink had dried the slides were washed 3 times, 5min each, with PBS. The tissue was permeabilized with 0.2% Triton-X100 (Sigma, #T8787-250ML) for 20min at RT, followed by incubation in blocking solution: 1×PBS containing 5% normal goat serum (Thermo, #16210-072), 1% BSA (Sigma, #A3912-100G), and 0.05% Tween-20 (Sigma, #P2287-100ML) for 1h at RT. All samples were incubated in primary antibodies (Table 1) in blocking solution overnight at 4°C. The following day slides were washed 4 times, 10min each, with 1×PBS containing 0.05% Tween-20 (PBST) and incubated for 90min with secondary antibody (Table 1) in blocking solution. Slides were washed 3 times, 10min each, in PBST and then incubated with bisbenzimide (Thermo-Fisher, #H3569) for 5min for nuclear counterstain. This was followed by 3 x 5min washes in 1×PBS, then coverslips were mounted on slides with Glycergel mounting medium (Agilent Dako, #C0563) and allowed to cure at RT O/N. The following day the slide edges were sealed with clear nail polish (EMS, #72180) and stored at 4.

**Table 1.** Antibodies.

| Antibodies Table 1 |  |  |  |  |
| --- | --- | --- | --- | --- |
| Antibody | Species | Dilution | Company | Cat# |
| **OCT 3/4** | Rt | 1 (250) | R & D Biosystems | #MAB1759 |
| **NANOG** | Rb | 1 (500) | Abcam | #ab21624 |
| **CTIP2/**<br>**BCL11B** | Rt | 1 (300) | Abcam | #ab18465 |
| **PAX6** | Rb | 1 (1000) | MBL | #PD022 |
| **SATB2** | M | 1 (100) | Abcam | #ab51502 |
| **TUBB3/TuJ1** | M | 1 (1000) | Covance | #MMS-435p |
| **TBR2/EOMES** | Mouse | 1 (400) | R&D | #MAB6166 |
| **HOPX** | Rb | 1 (500) | MilliporeSigma | #HPA030180 |
| **MKI67** | Mouse | 1 (1000) | BD biosciences | #550609 |
| **Anti-active** | Rb | 1 (1000) | BD | #559565 |
| **Caspase-3** |  |  |  |  |
| **SOX2** | Rb | 1 (1000) | MilliporeSigma | #AB5603 |
| **anti-Mouse IgG**<br>**Alexa Fluor 568** | Gt | 1 (500) | Invitrogen | #A-11031 |
| **Anti-Rabbit**<br>**IgG AlexaFluor**<br>**647** | Gt | 1 (500) | Invitrogen | #A-21245 |
| **Anti-Rat IgG**<br>**AlexFluor 488** | Gt | 1 (400) | Invitrogen | #A-11008 |

### Organoid size and fluorescence intensity quantification

To measure the size of organoids after vitrification and rewarming, individual organoids were placed into a low attachment U-Bottom 96-well plate and imaged using an Incucyte live imaging system (Sartorius) from days 0-10 post rewarming. Phase contrast images were taken at 10× magnification using the spheroid protocol. ICC images for quantifications were acquired on a DMI6000B epifluorescence microscope (Leica) equipped with a Hamamatsu Photonics Camera (#C4742-80-12AG). Representative images of individual organoids were taken under 20× magnification and were quantified for fluorescence intensity using Fiji based on a protocol described in Dang et al., 2021^25^. Briefly, 10X images were taken of each organoid containing as much of the organoid as possible. For nuclear proteins (KI67, Cleaved Caspase-3[Casp3], CTIP2/BCL11B, SATB2, SOX2, TBR2/EOMES, and PAX6), the percentage of area immunolabeled (% Area) was calculated by measuring the area that stained positive for each of the above markers that overlapped with bisbenzimide/DNA labeling. Bisbenzimide was used to create a region of interest (ROI) and any nuclear stain that fell within the ROI was counted as positive for one of the above markers. For cytoplasmic proteins (TUBB3/TuJ1, HOPX, GFAP) the whole organoid was set as the ROI. The relative intensity for immunolabeling was calculated for each cytoplasmic protein by dividing the mean intensity of immunoreactivity for the protein of interest by the mean fluorescence intensity of bisbenzimide within the same ROI. Representative images of the vitrification and control organoids were taken using an Andor BC43 spinning disk confocal microscope (Oxford Instruments).

### RNA Extraction and RT-qPCR

Total RNA was isolated with the RNeasy Mini Kit (Qiagen, #74104) according to the manufacturer’s protocol. cDNA was synthesized with the SuperScript™ III First-Strand Synthesis SuperMix kit (Invitrogen, #18080-051). The RNA integrity values (260/280 and 260/230) were equivalent between vitrified and control organoids. RT-qPCR was carried out with Power SYBR Green PCR Master Mix (Applied Biosystems, #4367659) using primers listed in Table 2. qPCR was performed on a Quantstudio 3 Real-Time PCR System (Thermofisher). Fold change was calculated using the 2^−ΔΔCt^ method: ΔΔCt = (Ct gene of interest [Vitri or control] – average Ct GAPDH) – (average ΔCT control sample gene of interest) eg. Vitrified ΔΔCt [PAX6] = (CT PAX6 – CT GAPDH) – (avg. ΔCT control PAX6).

**Table 2.**
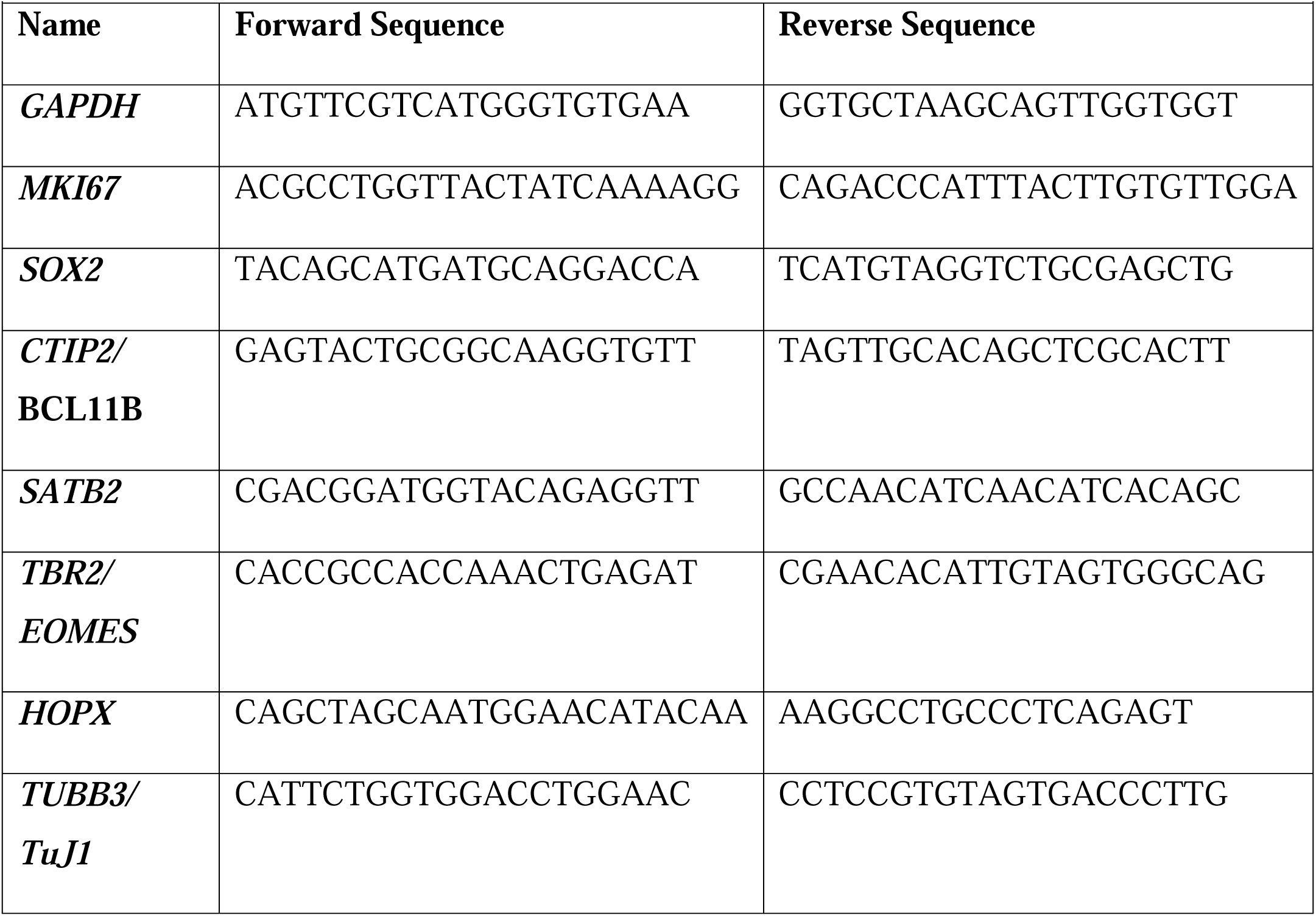
Primers.

### Electrophysiological Recordings

#### Multielectrode Array (MEA) Recordings

At day 90 post derivation (p.d.) SOSR-COs were plated on MEA plates to observe activity using a protocol adapted from Pan et al., 2024.^46^ Briefly, MEA plates (Axion Biosystems, #M384-tMEA-6W) were washed with 1 mL of 1% Tergazyme (Alconox, #1325) in ddH_2_O followed by 3 washes with ddH_2_O. 1mL of organoid culture medium was added to each well and placed in a 37°C incubator for 2 days. Media was aspirated and replaced with 0.1% polyethylenimine (PEI; Sigma, #P3143-100mL) in ddH_2_O and incubated at 37°C for 1h. Each well was rinsed 3X with ddH_2_O and allowed to air dry in the cell culture hood for 1h at RT with the lid off. Next, we made a 1:10 mixture of Geltrex in DMEM/F12 and added 50µL to the center of each well. The plates were left in the incubator O/N at 37°C. The Geltrex solution was then aspirated and any Geltrex covering the electrode contacts was sprayed away using DMEM/F12 and a p1000 pipette. Two organoids were transferred to each well of the MEA plate. The residual media was aspirated with a p200 tip, and the plate was placed in the 37°C incubator for 5min. The plate was then removed and a single drop of 100% Geltrex was applied to the top of each organoid using a p200 pipette tip. The plate was then placed back in the incubator for another 10min. Culture media (250µL) was added gradually over several hours. The next day the remaining 750µL of media was added and the plate was left in the 37°C incubator for 2 days. 3N media was gradually transitioned over the next week to BrainPhys™ media (STEMCELL Tech., #05790). Recordings were taken the day after media changes, which were performed twice per week using an Axion Maestro Original running Axis Software. Recordings were made using 200 Hz – 3kHz bandpass filtering and analyzed using the Axis Metrics Plotting Tool (Axion Biosystems).

#### Whole cell Patch Clamp Recordings

Day 90 SOSR-COs were transitioned from 3N media to BrainPhys™ media over 1 week and cultured until day 200. Organoid slice preparation and recordings were modified from Tidball et al., 2023^24^. Briefly SOSR-CO slices (∼300-400µm) were cut manually in ice-cold, oxygenated “slicing” solution saturated with 95% O_2_ /5% CO_2_ because the organoids were too small to cut with an oscillation tissue slicer. The slicing solution contained (in mM): 110 sucrose; 62.5 NaCl; 2.5 KCl; 6 MgCl_2_; 1.25 KH_2_PO_4_; 26 NaHCO_3_; 0.5 CaCl_2_ and 20 D-glucose (pH 7.35-7.4 when saturated with 95% O_2_ /5% CO2 at RT of 22-25°C). Slices were incubated in slicing solution for 20-30 min at RT and then in a 1:1 mixture of slicing solution and artificial cerebrospinal fluid (ACSF, containing (in mM): 125 NaCl; 2.5 KCl; 1 MgCl_2_; 1.25 KH_2_PO_4_; 26 NaHCO_3_; 2 CaCl_2_ and 20 D-glucose, (pH 7.35-7.4)) in a holding chamber aerated continuously with 95% O_2_ /5% CO_2_ at 35°C for 30min before being transferred to ACSF at RT. Individual slices were transferred to a recording chamber perfused (2-3 mL/min) with ACSF bubbled continuously with 95% O_2_ /5% CO_2_ at RT. Cells were visualized and selected using an E600FN upright microscope (Nikon) with a Namarski 40X water immersion objective. For action potential recordings, recording electrodes had a resistance of 4-6 MΩ when filled with a potassium gluconate (K-gluconate)-based pipette solution consisting of (in mM): 140, K-gluconate; 4, NaCl; 0.5, CaCl_2_; 10, HEPES; 5, EGTA, 5 phosphocreatine; 2, Mg-ATP and 0.4, GTP (pH 7.2-7.3 adjusted with KOH). Whole cell current-clamp recordings were used to measure APs. Repetitive firing was evoked by a series of 1500ms depolarizing currents varying from -60pA to 300pA in 10pA steps from the resting membrane potential.

Spontaneous firing of APs from their resting membrane potential was recorded using a gap-free mode. For recording sEPSCs in the same cells, recordings were switched to whole cell voltage-clamp recording mode. sEPSCs were recorded at a holding potential of -70mV in the presence of 10µM bicuculline in the external solution to block GABAergic synaptic responses. The signals were amplified using a Multiclamp700B amplifier (Molecular Devices), filtered at 2-4kHz and digitized at 20Hz for analysis. Data were acquired with a Digidata1440 digitizer and analyzed using pClamp11.3.

### Statistics

Data were organized and analyzed in Excel (Microsoft). All graphs and statistics were performed in Prism 10 (Graphad). Cell counts and qPCR results were tested for statistically significant differences using students *t-*Test. Size measurements, MEA recordings and input-output curves were analyzed using two-way-ANOVA. Error bars depict the standard error of the mean (SEM).

## Supplemental Information

**Supplemental Figure 1. Confirmation of pluripotency markers in reprogramed iPSC lines. A.** 79b iPSC derived from human foreskin fibroblast cells were stained for **Ai.** NANOG (green) **Aii.** OCT4 (green) and **Aiii.** SOX2 (green). All counterstained with Hoechst (blue). **B.** A2 cell line derived from foreskin fibroblast also demonstrates pluripotency markers: **Bi.** OCT4 (green), Nanog (red), and **Bii.** OCT4 (green) SOX2 (red). All counterstained with Hoechst (blue). Scale bar: 200µm.

**Supplemental Video 1.** Steps to perform vitrification of cortical hCOs.

**Supplemental Video 2.** Steps to warm hCOs after vitrification.

